# The best plant-guarding ants in extrafloral nectaried plants and myrmecophytes according to baiting tests

**DOI:** 10.1101/2023.03.03.530851

**Authors:** Leticia Silva Souza, Eduardo Soares Calixto, Saulo Santos Domingos, Alexandra Bächtold, Estevao Alves Silva

## Abstract

Extrafloral nectaried plants and myrmecophytes offer resources to ants that may engage in protective mutualisms. The role of different ant species in herbivore deterrence has long been analyzed by using herbivore baits, and ants are regarded as effective plant guards if they attack and/or remove the baits (mostly termites) from plants. Here, we conducted a comparative investigation on which ants display aggression toward baits, which ants are better plant guards, and which plants (extrafloral nectaried plants or myrmecophytes) are better defended by ants. Data from the literature revealed that baiting studies have been performed on 37 extrafloral nectaried plant species and 19 myrmecophytes, and have involved over 30 genera of ants. Extrafloral nectaried plants and myrmecophytes rely on specific ant fauna to defend them from herbivores. In extrafloral nectaried plants, *Camponotus* and *Crematogaster* were regarded as the best plant protectors, as they attacked baits in nearly all plants. In myrmecophytes, *Azteca, Pheidole* and *Pseudomyrmex* were the most important bait attackers. Myrmecophytes were better protected by ants, as all baits were attacked; in extrafloral nectaried plants, some ants failed to attack the baits. Plants can be patrolled by several different ants, but there is a core of ants that excel in protection, and this varies according to plant type (extrafloral nectaried plants and myrmecophytes). With this knowledge, it may be possible to label different ants as effective plant guards, to anticipate their effects on plant performance and even to understand their potential role as biological control agents.

## 1. Introduction

Mutualistic interactions between plants and ants are shaped by the provision of shelter and/or food resources by plants (e.g., nesting sites, extrafloral nectar) in exchange for ecological services by ants, who act as bodyguards by defending the plants from herbivores (Cagnolo and Tavella, 2015; Oliveira et al., 2021). Plants that offer resources to ants are represented by myrmecophytes and extrafloral nectary (EFN) bearing plants. Myrmecophytes usually provide ants with shelter, domatia and/or food resources (Dejean et al., 2017), while EFN plants usually provide only extrafloral nectar, a carbohydrate-rich solution (Nogueira et al., 2015). The interaction between ants and myrmecophytes is often obligate and involves a few ant species associated with each plant species while EFN plants sustain a facultative interaction with ants, meaning non-specialization between both parties (Blüthgen et al., 2007; Fiala et al., 1999). This gradient between facultative and obligatory interactions can therefore impact the results of the protection mutualism (Fiala et al., 1994).

Given the richness of ants found in plants, it is not surprising that some are neither aggressive nor exert a role as bodyguards (Leal et al., 2006). This results in two contrasting scenarios. On the one hand, there is a core of ants that maintains a beneficial relationship with plants; on the other hand, there are ants that benefit from plant resources but do not offer a countermeasure (Nogueira et al., 2012). In addition, in some cases, ants may even negatively influence plant performance or fail to deter herbivores (Melati and Leal, 2018). Evaluating ant protection effectiveness, therefore, is essential to better understand which ants establish mutual interactions with plants, potentially reducing the damage inflicted by herbivores and increasing the reproductive success of plants (Raupp et al., 2020).

To investigate ant protection effectiveness, scientists often use herbivore baits to analyze ant aggressiveness (Calixto et al., 2021). Baiting tests start with the introduction of a bait, e.g., termites or caterpillars, on the plant, and ant behavior is recorded within a given period (Oliveira et al., 2021). Protection is considered successful whenever ants attack or remove the bait (Fagundes et al., 2017). This procedure has provided important insights to explain why some ants perform better than others in herbivore deterrence and plant protection (Lach and Hoffmann, 2011). For instance, *Pseudomyrmex* ants were labeled as effective plant guards because they attacked more bait and were more alert than other ant species that co-occurred on the same plants (Oliveira et al., 1987). However, to our knowledge, there is no extensive comparative investigation on which ants display aggressive behavior toward herbivore baits, which ants are better plant guards, and which plants (myrmecophytes or EFN plants) are better defended by ants. A close examination of ant aggressiveness might permit us to label different ants as effective plant guards, to anticipate their effects on plant performance, and to understand their potential role as biological control agents (Kenne et al., 2000; Schatz et al., 1997).

In this study, we examined the literature regarding baiting tests, and we retrieved information on the ant species examined, the ant behavior (attack), and the types of study plants (EFN plants and myrmecophytes) where tests were performed. With this data, we investigated *(i)* whether some ants were bait-tested more than others and *(ii)* if ant attack was related with plant type. This might give us an indication of which ants are more common in myrmecophilous plants and whether and where they attack most baits (Fagundes et al., 2017; Leal et al., 2006).

Furthermore, in line with the degree of ant protection exerted by each ant that foraged on a plant population (Fagundes et al., 2017; Tillberg, 2004), we *(iii)* investigated the ants that were regarded as better protectors in EFN plants and myrmecophytes. Within the bait-tested ant community, some ants may excel and be regarded as better bodyguards; thus we hypothesized that a core of ants would outstand in terms of attacking baits (Cruz et al., 2018; Leal et al., 2006).

To conclude we *(iv)* analyzed which EFN-plants and myrmecophytes were defended by only one ant genus, named here as sole defenders (the reasons for using ant genera are clarified in the Methods). We expected that more species of myrmecophytes would rely on sole defenders for protection because of the close and specific association between ants and myrmecophytes in comparison with EFN plants, where more ants are assumed to interact with plants (Blüthgen et al., 2007; Cagnolo and Tavella, 2015).

## 2. Materials and Methods

### 2.1. Literature search

The search for literature of baiting tests was made using the words “ants, baits, termites, extrafloral nectar, myrmecophytes, myrmecophily, ant aggressiveness” and variations. The Google Scholar (https://scholar.google.com.br/) was the only internet searching tool used, as it reunites literature from all publishers and usually permits the direct download of the paper, either from the publishers’ website, or from other platforms such as the Research Gate or the personal webpage of scientists. Our search covered papers published as early as 1981 (first study that examined ant attack toward baits) until December 2021. After a thorough search, we came across dozens of studies which were downloaded and read. During the reading of papers, we also scrutinized the references list to gather additional literature. We also came in touch with scientist to request studies that were not fully available on the internet.

### 2.2. Criteria to standardize the data

To standardize the research, only the papers that fulfilled the following criteria were included in our analyses: (i) published papers, as this literature type undergoes peer review; (ii) papers in English, for permitting the checking of data by anyone in the scientific community; (iii) studies that covered arboreal ants; (iv) studies conducted in the field; (v) papers in which the methods clearly specified the use of animal baits, such as termites and other insects; (vi) papers with taxonomic identification (at least genus) of the study ants and plants; (vii) studies in which the information of the ants associated with each plant could be retrieved; (viii) papers that indicated which ants engaged in aggressive behavior toward baits. On some occasions authors showed only a subset of ants that demonstrated aggressive behavior; according to these authors, these ants were regarded as the most important in the community. This data was included in our research.

### 2.3. Data collection

Each paper included in the analyses was named and number labeled. The following information was retrieved and systematically organized: the country where the fieldwork took place, the taxonomic information of ants and plants, the baits used (e.g., termite, caterpillar, grasshopper), the type of study plant (‘EFN’ – plants that produce extrafloral nectar – Del-Claro et al. 2016; and ‘myrmecophytic’ – plants that shelter ants and may have or not have food resources – Dejean et al. 2018), and whether ants attacked the baits.

An ‘attack’ consists of the aggression of ants, which varies from species (biting, stinging, spraying liquids), but we also considered the molestation of ants as attack, as this is also an antiherbivore mechanism (Alves-Silva and Del-Claro, 2014; Fuente and Marquis, 1999). Furthermore, for small ants, it is difficult to determine if they bite the baits; in some instances, the authors did not explicitly mention that ants really bit the baits, but this was implied in the study (Leal et al., 2006; Vidal et al., 2016).

The repertoire of ant behavior toward baits is extensive (Dejean et al., 2009). For instance, recruiting and bait removal from plants might also provide evidence of plant protection. Recruitment involves the arrival of other nestmates at the place where the bait is being attacked (Pacelhe et al., 2019). Removal consists of removing the bait from the plant by either taking it to the nest, throwing it away, killing it or making it abandon the plant (Alves-Silva and Del-Claro, 2016; Vantaux et al., 2007). Despite the importance of both behaviors in herbivore deterrence, scientists rarely provide such information; in addition, these behaviors are not subjected to tests (such as ant attacks toward baits), but rather sporadically noted in the field. In fact, studies rarely specify whether ants display recruitment and removal of baits. This may derail detailed comparisons and even bias conclusions. However, the attack (or not) of ants on baits is always specified. Thus, to keep the data as appropriate as possible to allow statistical comparisons and to increase our understanding of ant behavior, we examined only ant attacks.

### 2.4. Ants who engage in aggressive behavior

Our data cover ant genera rather than species, and this is justified. Ants were identified to species level in approximately 2/3 of baiting tests, but this number varied according to plant types; for instance, only 58% of the ants in myrmecophytes were identified to species level, especially due to the difficulty of dealing with small ants, such as *Azteca* and *Crematogaster*. Thus, the use of ant genera is substantiated by the difficulty of some authors in identifying some ants at the species level (Koch et al., 2016). Many ant species were observed in the field and classified into genera but were not collected and subjected to further taxonomic identification. In some cases, new species were classified as ‘sp. nov’ (and variations) and no specific name was provided.

By using ant genera, we also avoided considering ant species that were misidentified to the species level or species that belong to a group of species (e.g., *Camponotus* aff. *blandus, Pseudomyrmex* gr. *pallidus*). We also noted misspelling in specific names, which indicates minimal taxonomic attention to the species level in some cases. To conclude, the approach of focusing on ant genera to classify ants as effective plant guards is valid when we consider that arboreal species in many genera present conservative behavior when it comes to foraging, patrolling and aggressiveness toward baits (Bächtold and Del-Claro, 2018; Fagundes et al., 2017; Oliveira et al., 1987).

### 2.5. Data analyses

We found 56 papers that fulfilled our study criteria and we managed to retrieve 344 records of ants that were bait-tested. Exploratory analyses revealed the presence of 33 ant genera, however the most frequent ants belonged to nine genera, namely *Camponotus, Crematogaster, Pseudomyrmex, Pheidole, Cephalotes, Azteca, Ectatomma, Solenopsis* and *Brachymyrmex* (Supplementary material 1), which together were present in 85% of papers (*n* = 48), 85% of baiting tests (*n* = 292) and at least in one plant type (EFN and/or myrmecophytes). Our analyses will cover the data of these nine genera of ants only (*n* = 292, the statistical unit of our study), as it permits the use of statistical analyses and comparisons.

### 2.6. Statistical analyses

Goodness of fit chi-squared tests were used to check if one ant genus was tested more than others; we compared the frequency of each ant in baiting tests, in the occurrence of plant species, plant families, plant types (EFN and myrmecophytes) and in the papers that bait-tested ants (objective *i*). Two-way Anova tests were conducted to analyze if ant attacks were related with the ant genera and plant-types (objective *ii*).

To investigate the importance of each ant as plant-guard (objective *iii*), we built ecological networks with pairwise interactions between ants and the plants where they attacked baits. The ecological network was defined by the connections between plants and ants, and it is considered a holistic approach to investigate ecological communities (e.g., pollination, predator-prey interactions, mutualisms) and how species interact with their partners (Trøjelsgaard and Olesen, 2016). Both ants and plants were regarded as nodes, and the interactions between them were labelled as links (Guimarães, 2020).

We built adjacency matrices where values of one or zero were assigned to cells with or without interactions between ants and plant species, respectively. According to Palacio et al., (2016 and references therein) binary data (presence and absence of interactions between species) are appropriate to calculate centrality indices (Mello et al., 2015) and results do not differ in comparison to weighted data (i.e., how often a species *i* interact with species *j*); in addition binary data may identify central nodes as good as weighted data does. With the organization of the matrix and further analyses of data we could see the plants defended by each ant, possible overlaps of different ants in a single plant species, and which plants relied on only one ant genus for defense (sole defenders). A total of two matrices were built, one for EFN-plants and other for myrmecophytes.

We calculated the degree, which shows the number of plant species where ants were observed to attack baits (Cagnolo and Tavella, 2015) and we also estimated the centrality values for ants in each network. Centrality indices have been incorporated in ecological studies to determine the important species in communities (Farine and Whitehead, 2015; Genrich et al., 2017; Martín González et al., 2010; Poulin et al., 2013). Collectively, the centrality indexes not only consider the quantity of interactions between parties (degree distribution), but also the indirect connections of a species with the whole plant community (Delmas et al. 2019; Silva et al. 2020). Thus, a value of the relative importance of a species in a community is provided (Borrett, 2013; Martín González et al., 2010).

In our study we used the betweenness (BC) and the eigenvalue (EC) centrality indexes. The first gives more weight to ants that have higher degree (i.e., ants that attack baits in many plant species) and if removed cause the rupture of the network (Olesen et al., 2007). In this context, ants with high BC are better plant-guards, because they attack baits in several plant species, may act alone in a few plant species, and if removed may leave plants either unattended or patrolled by so-called less aggressive ants (by less aggressive we mean those ants that occur in a few plant species and are thus not frequent in the plant community) (Sazima et al., 2010; Trøjelsgaard and Olesen, 2016).

The EC gives weight to overlaps between ant species in plants. Ants that occur in many plant species will have higher EC, however, the less frequent ants that co-occur with the most frequent ants will also have higher EC (Borgatti, 2005; Morand et al., 2020; Silva et al., 2020). This index indicates the contribution of ants to the plant community and also shows the specialization, as ants with the lowest EC will be rare in the community and will not share plants with frequent ants.

The metrics used in plant-animal networks are almost limitless (Delmas et al., 2019; Guimarães, 2020), but we used only those that were suitable to address our objectives. For instance, the closeness centrality (the occurrence of a species *i* in the same niches/plants of other species) is also used in some studies (Mello et al., 2015), but it positively relates with degree and BC (Sazima et al., 2010). In addition, we calculated the metrics for ants only; the species of plants were not incorporated into the metrics as we were interested in the functional groups they belonged to (i.e., EFN-plants or myrmecophytes), rather than on particular plant species. Our objectives are in accordance with the definition of microscopic metrics of the network as we focused on the importance of species (in our case, the different ants) rather than the overall network structure (Trøjelsgaard and Olesen, 2016). All centrality indices were calculated with the *igraph* package in R statistical software (Csardi and Nepusz, 2006).

To conclude, we investigated which plant species sustained only one ant genus for protection. Most plants were patrolled by more than one ant genus, whereas others relied on only one genus of ants as a plant-guard. We separated the ants that attacked baits on only one plant species and these ants were regarded as sole protectors. We then used the Fisher’s exact test to examine which plant type (EFN-plants and myrmecophytes) had more sole defending ants (objective *iv*).

We are aware that some interactions in the plant-ant network may be forbidden links, i.e., interactions that are likely to not occur (Kiziridis et al., 2020). Given the spatial scale of our data (information retrieved from 56 papers that bait-tested ants in eight countries), the forbidden links were expected to occur. We are fully aware of the issues related with forbidden links (Jordano, 2016). Nonetheless, the purpose of our network was to show that among the pool of ants, some genera are regarded as more important than others in plant defense; and if that particular ant (genera) is removed, the plant can be occupied by ants with similar aggressiveness, or not. In addition, the most important ants in our work are widespread in the world and can overlap in several plant species (shown in Results). To conclude we did not consider particular plant species, but rather if plants either belonged to the EFN or the myrmecophyte group.

We provide an overview of the characteristics of baiting studies, such as where investigations were performed, the bait types used and the study plants, but without statistical tests. Supplementary materials bring figures with more detail of the data.

All the analyses and figures were made in R statistical software version 4.1.3.

## 3. Results

### 3.1. Overview of baiting studies

The baiting studies which were performed with the most frequent ants (*Camponotus, Crematogaster, Pseudomyrmex, Pheidole, Azteca, Ectatomma, Solenopsis, Cephalotes* and *Brachymyrmex*) were conducted in eight countries, half of them in Brazil (50% of the total, *n* = 24 studies); 60% of the baiting tests were also made in Brazil (*n* = 174). Termites were used more often (81% of tests) than the other bait types, such as caterpillars, beetles, orthopterans, flies, and hemipterans (Supplementary material 2). The plants were represented by 56 species in 37 genera and 26 families, especially Fabaceae (30% of plants species, *n* = 17 species) (Supplementary material 3). A total of 37 EFN-plant species and 19 myrmecophytes were studied (Supplementary material 4).

### 3.2. Ants in baiting tests

Both *Camponotus* and *Crematogaster* were significantly more frequent in baiting tests, plant species, plant families, and in the number of studies that had these ants under investigation (Table 1). Their frequency in EFN-plants was also significantly higher in comparison to the other ants examined (Table 1). In myrmecophytes, there was no difference in the frequency of ants.

**Table 1.** Frequency data of the most common ants in baiting studies. Superscript lowercase letters indicate statistically significant differences, according to chi-squared tests.

| Ants | Baiting tests | Plant species | Plant families | EFN plants | Myrmecophytes | Papers |
| --- | --- | --- | --- | --- | --- | --- |
| *Camponotus* | 98<sup>a</sup> | 37<sup>a</sup> | 18 | 33<sup>a</sup> | 4 | 29<sup>a</sup> |
| *Crematogaster* | 47<sup>b</sup> | 30<sup>ab</sup> | 15 | 25<sup>ab</sup> | 5 | 23<sup>ab</sup> |
| *Pseudomyrmex* | 41<sup>c</sup> | 23<sup>ac</sup> | 8 | 18<sup>a</sup> | 5 | 14<sup>a</sup> |
| *Cephalotes* | 23<sup>d</sup> | 16<sup>b</sup> | 8 | 14<sup>b</sup> | 2 | 12<sup>ab</sup> |
| *Pheidole* | 23<sup>d</sup> | 14<sup>c</sup> | 6 | 9<sup>c</sup> | 5 | 12<sup>ab</sup> |
| *Azteca* | 18<sup>d</sup> | 9<sup>c</sup> | 6 | 2<sup>c</sup> | 7 | 12<sup>ab</sup> |
| *Ectatomma* | 16<sup>d</sup> | 10<sup>c</sup> | 5 | 8<sup>c</sup> | 2 | 8<sup>d</sup> |
| *Solenopsis* | 14<sup>d</sup> | 10<sup>c</sup> | 7 | 9<sup>c</sup> | 1 | 8<sup>d</sup> |
| *Brachymyrmex* | 12<sup>d</sup> | 10<sup>c</sup> | 6 | 9<sup>c</sup> | 1 | 10<sup>b</sup> |
| Chi-squared | $\chi^2 = 184.88$<br>$P < 0.0001$ | $\chi^2 = 46.52$<br>$P < 0.0001$ | $\chi^2 = 18.86$<br>$P < 0.05$ | $\chi^2 = 53.35$<br>$P < 0.0001$ | $\chi^2 = 10.18$<br>$P > 0.05$ | $\chi^2 = 28.51$<br>$P < 0.001$ |

### 3.3. Ant attacks

EFN-plants were studied more often than myrmecophytes (*n* = 242 and 50 tests, respectively) (Figure 1a). The *Camponotus* was by far more frequent in EFN-plants than in myrmecophytes; *Azteca* was the only ant that was more frequent in myrmecophytes. In myrmecophytes, all baits were attacked; in EFN-plants, attacks were successful in 88% of tests (*n* = 212 tests). All ants attacked baits in both plant types (Figure 1b). Ant attacks were significantly related with plant types, but not ant genera (Table 2).

**Fig. 1.**
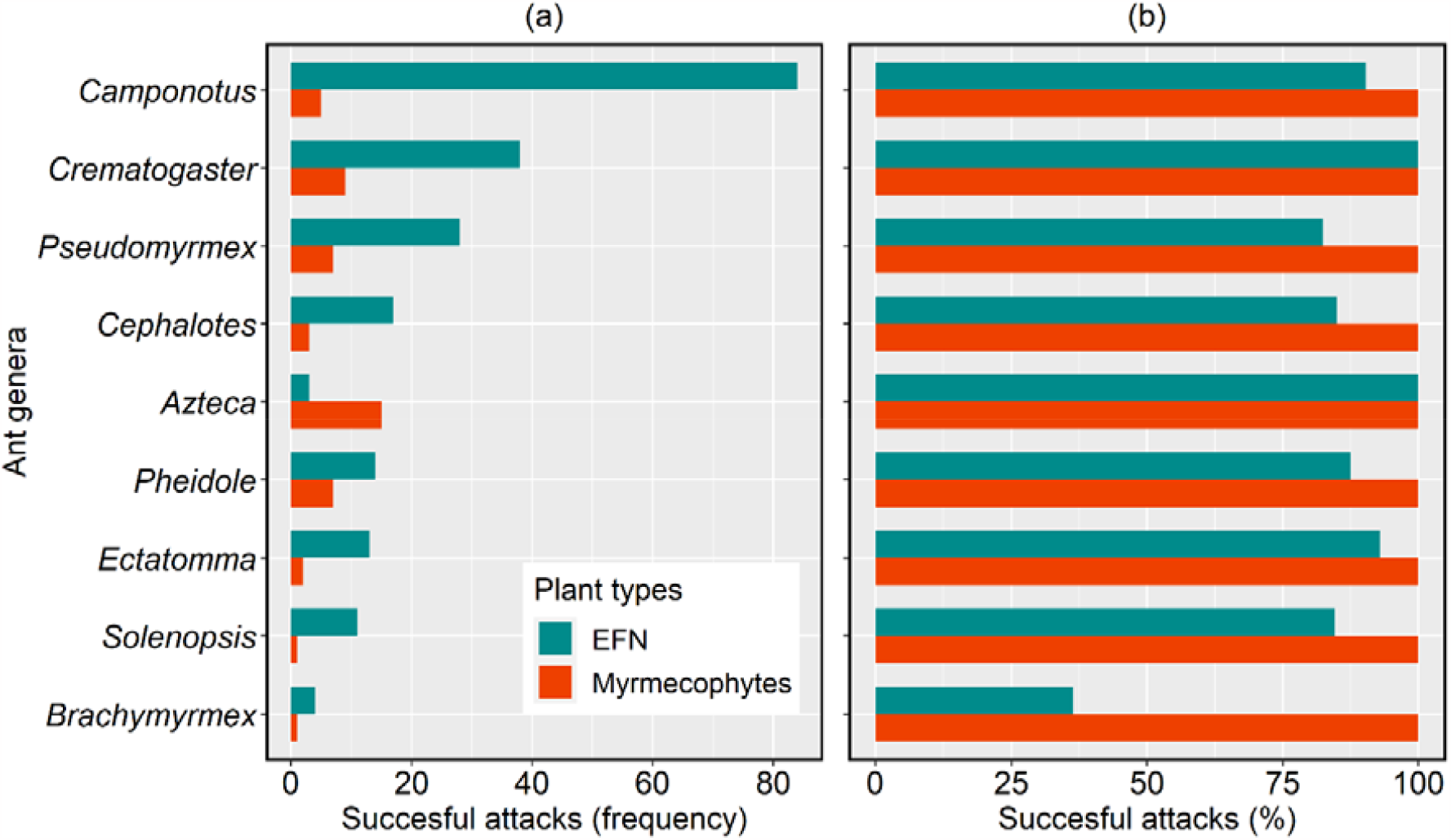
Ant genera that engaged in bait attack (frequency – a; successful attacks – b). EFN-plants sustained more baiting tests, with *Camponotus* and *Crematogaster* being frequently bait-tested. Myrmecophytes had 100% of baits attacked by ants.

**Table 2.**
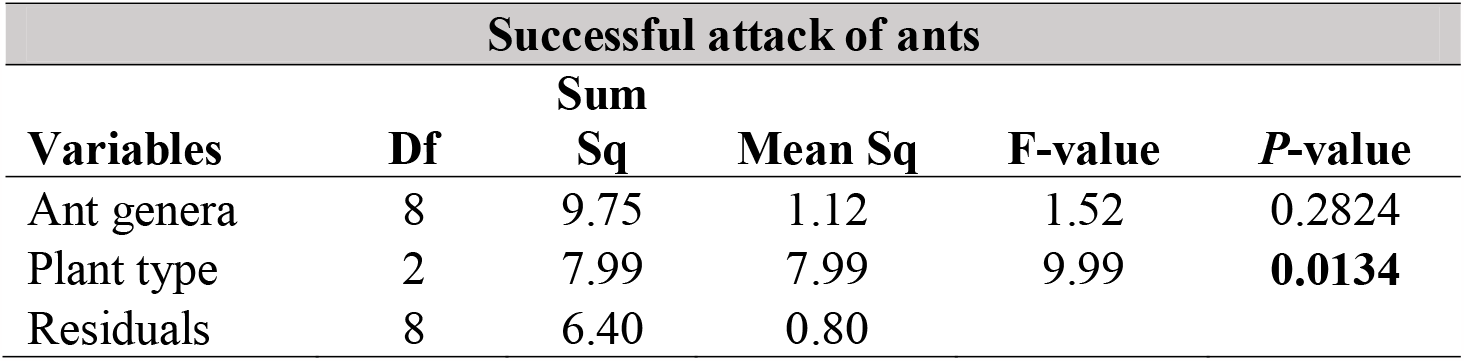
Anova indicating whether the frequency of attacks (log-transformed data) was influenced by the ant genera or plant type (i.e., EFN and myrmecophytes).

### 3.4. Important ants in the network

*Camponotus* and *Crematogaster* attacked baits in most plants within the EFN-plant community and had the highest degree (Figures 2a and 3a). These ants had also the highest BC and EC, indicating both that they were central to the network stability and they were connected to plants with other ants (Figures 2b, 3a, 3b). The BC was almost negligible for *Azteca, Brachymyrmex, Solenopsis* and *Pheidole*, indicating null to low importance in attacking baits in the EFN plant community; their EC shows that they shared a few plants with the most influential nodes in the EFN network (Figures 2c and 3b).

**Fig. 2.**
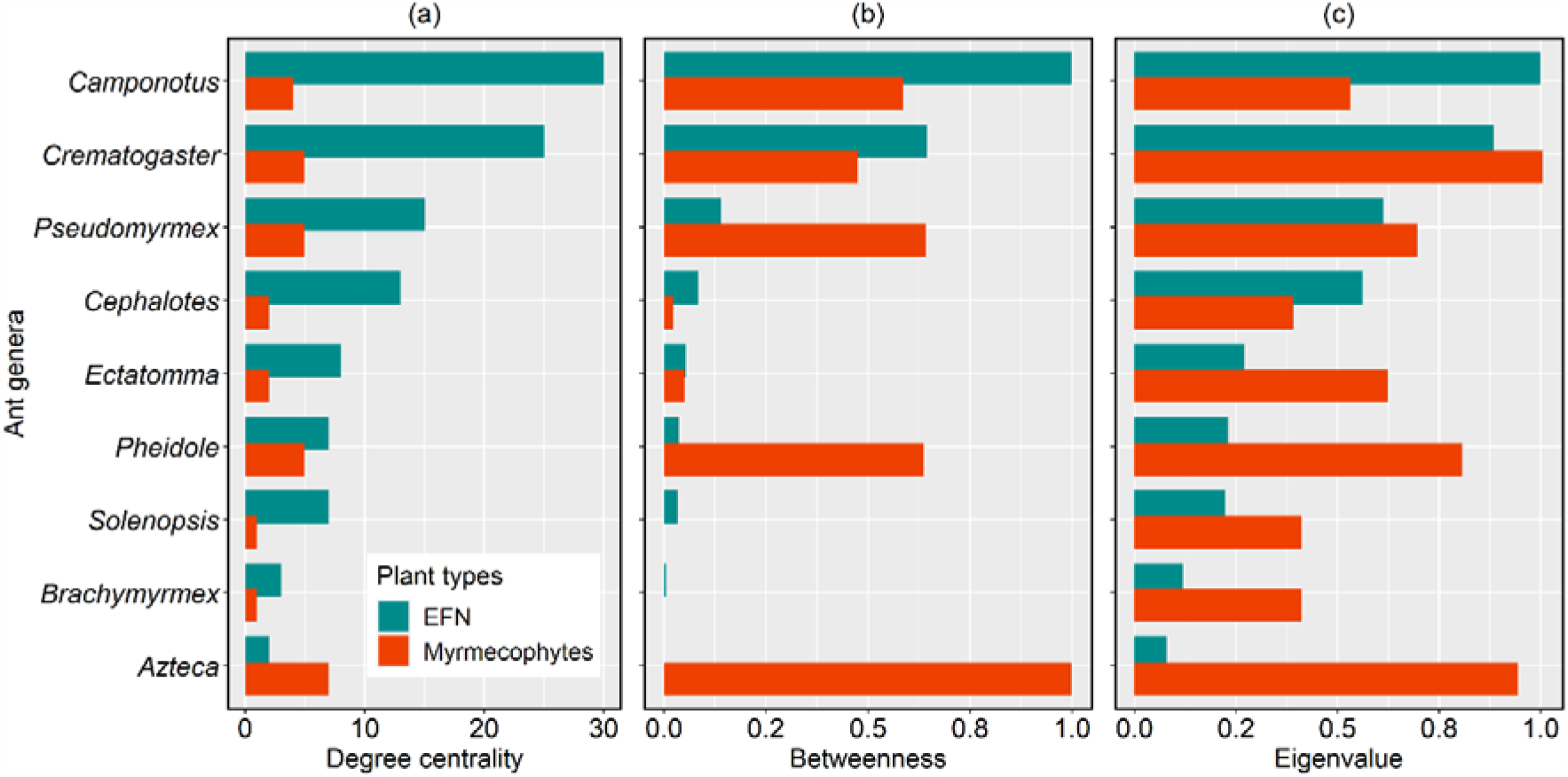
Measures of the relative importance of ants according to baiting tests. (a) Degree centrality which shows the number plants where each ant was bait-tested; (b) betweenness centrality (BC), that indicates ants that were central to the stability of the network; (c) eigenvalue centrality (EC) that measures co-occurrence of ants in the plant community. Values of BC and EC are relativized (0 to 1, lowest and highest value for a species, respectively) to permit a better visual comparison among plant communities.

The *Azteca* had the lowest degree, BC and EC in EFN plants, but they stood out in myrmecophytes with the highest degree, maximum BC and the second highest EC (Figures 2 and 3). This demonstrates how important were the *Azteca* in herbivore deterrence in myrmecophytes. The *Azteca* were followed by *Pheidole, Pseudomyrmex* and *Crematogaster* (Figures 2 and 3). Other ants (*Solenopsis, Brachymyrmex, Cephalotes* and *Ectatomma*) occurred in few myrmecophytes and their occurrence overlapped with the most aggressive/frequent ants; thus their BC and EC were low in myrmecophytes (Figures 2 and 3).

**Fig. 3.**
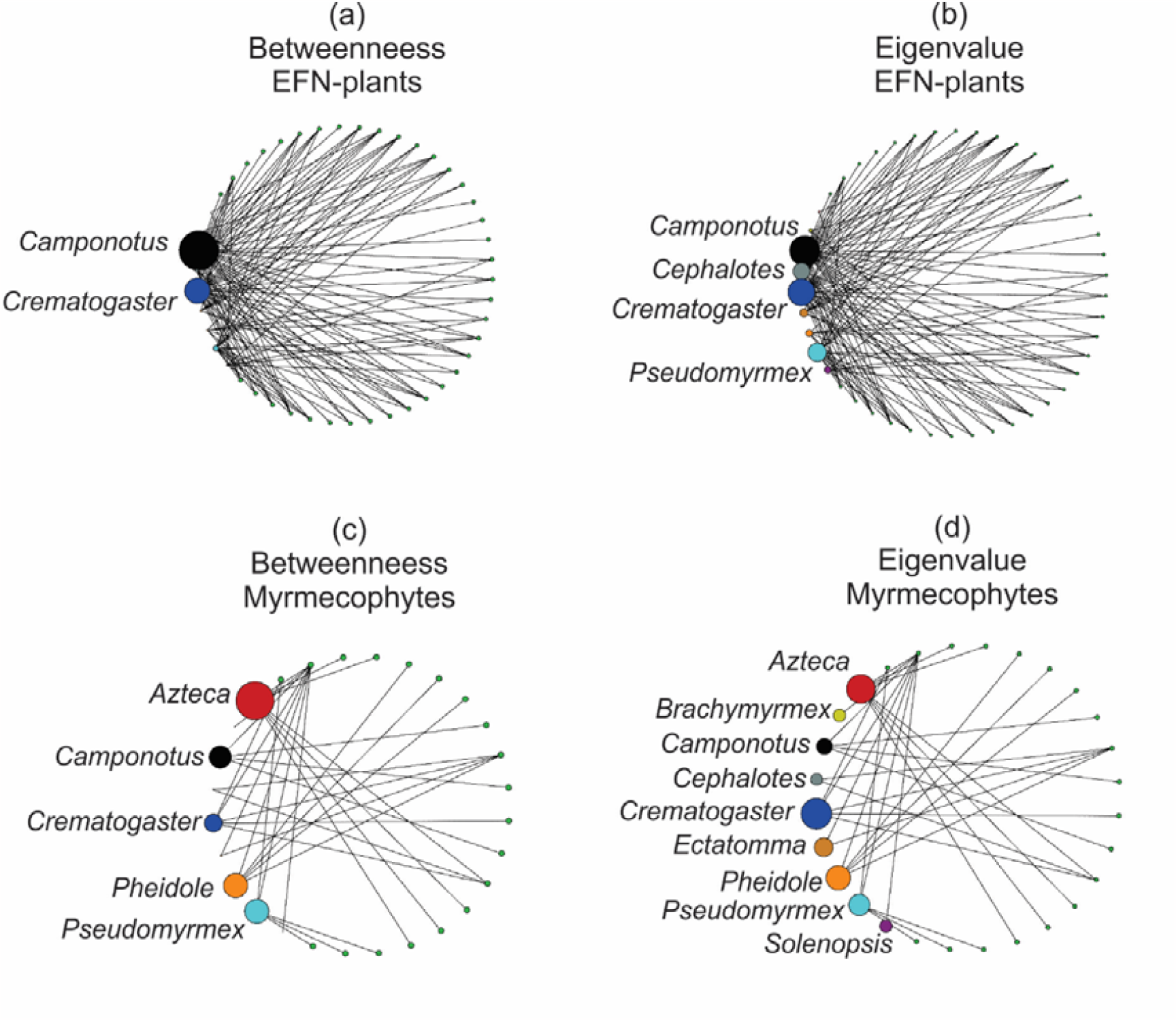
Ants that engaged in aggressive behavior to baits in (a, b) plants with extrafloral nectaries and (c, d) myrmecophytes. The small green circles represent the different plant species (names not shown). The size of circles indicates the relative importance of each ant according to the betweenness (BC) and eigenvalue (EC) centrality indexes.

### 3.5. Sole defenders

Out of the plants examined, nine EFN-plants and 15 myrmecophytes relied on only one ant genus for defense (Fisher’s exact test, *p* < 0.01). The *Azteca, Pheidole* and *Pseudomyrmex* were the sole defenders of myrmecophytes (*n* = 5, 3 and 3 plants, respectively), while the *Camponotus* and *Crematogaster* were the sole defenders of some myrmecophytes and EFN-plants (Figure 4).

**Fig. 4.**
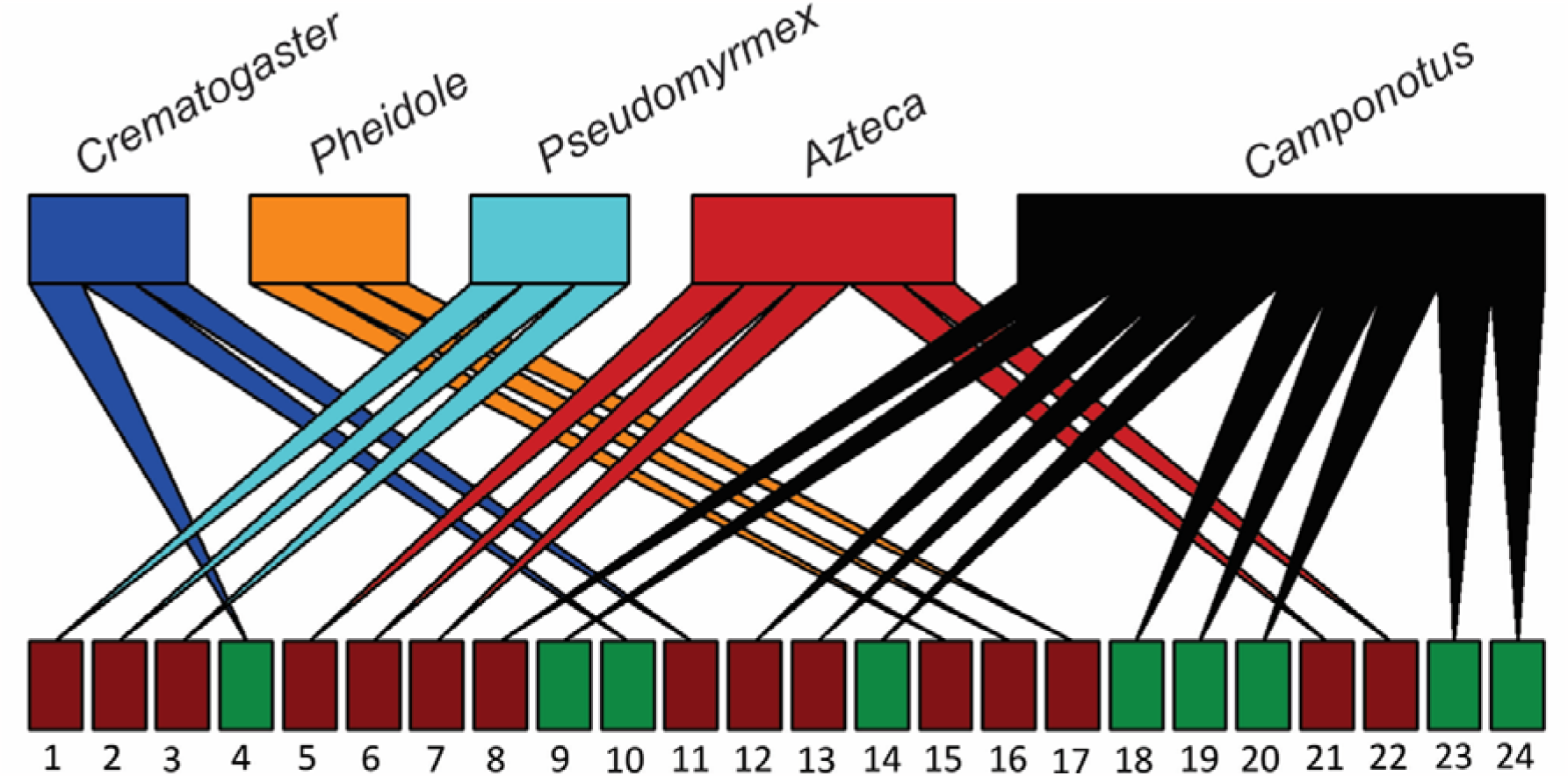
Bipartite network showing the ants that were sole defenders of EFN-plants (green) and myrmecophytes (dark red). Plant names: 1 *Acacia collinsii*, 2 *Acacia cornigera*, 3 *Acacia hindsii*, 4 *Cassia javanica*, 5 *Cecropia obtusa*, 6 *Cecropia obtusifolia*, 7 *Cecropia pachystachya*, 8 *Korthalsia furtadoana*, 9 *Lafoensia pacari*, 10 *Leea aculeata*, 11 *Macaranga banca*, 12 *Macaranga puncticulata*, 13 *Nepenthes bicalcarata*, 14 *Ouratea spectabilis*, 15 *Piper fimbriulatum*, 16 *Piper sagittifolium*, 17 *Piper* sp., 18 *Qualea multiflora*, 19 *Stachytarpheta glabra*, 20 *Stryphnodendron polyphyllum*, 21 *Tetrathylacium macrophyllum*, 22 *Tococa formicaria*, 23 *Triumfetta semitriloba*, 24 *Vachellia constricta*.

## 4. Discussion

### 4.1. Overview

Each plant type had a core of the most important ants in terms of bait attack, thus corroborating our hypothesis. With all results combined, *Camponotus* and *Crematogaster* excelled in EFN plants, and *Azteca, Pheidole* and *Pseudomyrmex* were the main bait attackers in myrmecophytes. The importance of these five ants was highlighted by their high centrality indices, indicating that they are frequent bait attackers in the plant community and that their absence could make plants unattended or attended by less frequent ants. These ants were also the sole defenders of some plants, indicating that without them, plants might rely on no ants as partners.

### 4.2. Core of the most important ants

*Camponotus* and *Crematogaster* were the most bait-tested ants and had the highest degree in many cases, except in myrmecophytes, where *Azteca* was dominant. In plant–ant relationships, there is usually a core of ants that dominate the interactions with plants; thus, it is expected that the most frequent ants receive higher importance in networks (Dáttilo et al., 2013; Lange et al., 2013; Silva et al., 2020). The centrality indices used in our study (betweenness and eigenvalue) not only considered the degree (how many plants ants interact with) but also the specific connections between ants and plants (Mello et al., 2015; Silva et al., 2020). More weight was given to ants that, whenever removed, might disrupt the network, leave plants unattended or be attended by less frequent ants (Genrich et al., 2017). Thus, species degree may not be related to other centrality values in some cases. For instance, both *Pseudomyrmex* and *Crematogaster* occurred in five myrmecophytes, but the former had higher BC; thus, *Pseudomyrmex* was more important to myrmecophytes than *Crematogaster*. This occurred both because *Pseudomyrmex* was present in myrmecophytes that had no other ants (sole defender of three plants), and because these ants shared only two plants with other ants. In contrast, *Crematogaster* attacked baits in plants that supported four other ants; that is why the EC was higher for *Crematogaster*, in comparison to *Pseudomyrmex*. Thus, in the absence of *Crematogaster*, plants can be patrolled by more ants than plants deprived of *Pseudomyrmex*. Interpretation of the different centrality indices (see Delmas et al., 2019; Palacio et al., 2016; Poulin et al., 2013) is paramount to understand the relative importance of each node for the stability of the network.

*Azteca, Pheidole* and *Pseudomyrmex* had the highest degree and BC values in myrmecophytes. Due to their small size, these ants can shelter and breed inside domatia and rapidly respond to disturbances in plants (Goheen and Palmer, 2010; Pringle et al., 2014). The specificity of this interaction makes *Azteca, Pheidole* and *Pseudomyrmex* the most important in myrmecophytes (Oliveira et al., 1987; Schmidt and Dejean, 2018). Since they are commonplace in myrmecophytes, they are likely to attack more baits. In contrast, EFN plants sustain a non-specific community of ants that do not shelter on plants but rather nest on the soil and forage on plants for food resources (Blüthgen et al., 2006). Even so, the ants in genera *Camponotus* and *Crematogaster* are constant in samplings (Burger et al., 2021; Koptur, 1984; Oliveira et al., 2021); these ants were more bait-tested, attacked more baits and received high BC values.

### 4.3. Effective plant guards?

Ecologists tend to be cautious in classifying a given ant as an effective plant guard because the aggressive behavior and protection efficiency of each ant species should be seen as a continuous rather than ordinal scale (Ness, 2006). In addition, the benefits ants provide to plants depend on two undissociated factors: quality and quantity. The former refers to the type of benefits received by plants, which in our case is bait attack, a proxy for herbivore deterrence (Calixto et al., 2021; Oliveira et al., 1987; Philpott et al., 2008) and the latter is the frequency of interactions between ants and plants (Koch et al., 2018; Ness, 2006). The combination of attack and frequency permitted us to draw a hierarchy of the importance of ants, with *Camponotus, Crematogaster* (EFN plants), *Azteca, Pheidole* and *Pseudomyrmex* (myrmecophytes) as the top plant defenders. In fact, some plants were observed to depend upon these ants only (the sole defenders), which is evidence of their importance as bodyguards in the plant community.

Intrinsic ant behavior and frequency in the plant community might be a good indication to label ants as effective plant guards. For instance, *Ectatomma* ants are aggressive and possess a privileged weaponry (body size, large mandibles and sting) that subdues several types of herbivores (Bächtold and Alves-Silva, 2013; Del-Claro and Marquis, 2015; Robbins, 1991; Schatz et al., 1997), but they are not regularly found on plants (Bächtold et al. 2016). In contrast, *Camponotus* and *Azteca* are shorter than *Ectatomma* and do not possess large mandibles or stings, but they consistently forage on many plant species (Blüthgen et al., 2000; Lange et al., 2019; Monique et al., 2022). Thus, they might exert a relatively higher role in herbivore deterrence in the plant community than infrequent ants (Calixto et al., 2021; Frederickson, 2005).

One may assume that the classification of ants as effective plant guards is premature. In fact, ideally all ants should be tested equally and then scientists could compare the attack rates among the ant community. Nonetheless, we do not believe that the different sample size of bait-tested ants is a drawback, as our dataset comprises a microcosm of plant–ant studies. The frequency of bait-tested ants in the literature reflects the occurrence of these ants in the field (Del-Claro and Marquis, 2015; Fagundes et al., 2017; Lange et al., 2013). *Camponotus* and *Crematogaster* are the most sampled species in the EFN plant community (Ribeiro et al., 2018) and the *Azteca*–myrmecophyte system is the most studied (Dejean et al., 2009). Our data covers papers from as far back as 40 years ago, and *Camponotus, Crematogaster* and *Azteca* have been consistently sampled ever since, indicating both that these ants are widespread in the field and that plant–ant interactions have temporal stability, with the core ant species occupying the same plants over time (e.g., *Camponotus* – Oliveira 1997; Koch et al. 2016; *Crematogaster* – Guimarães et al. 2006; Franco and Cogni 2013; *Azteca* – Tillberg 2004; Pringle et al. 2014). According to Díaz-Castelazo et al. (2010), generalist ants maintain the stability of the network and remain over time, even with the inclusion of other ants in the system. In this context, the results found here reflect what we actually see in the field in terms of plant occupation by ants.

## 5. Conclusions

The importance of ants, such as *Camponotus, Crematogaster* (EFN plants), *Azteca, Pseudomyrmex* and *Pheidole* (myrmecophytes) for herbivore deterrence is highlighted by their frequency on plants, their intrinsic aggressive behavior, the bait attack and their sole occurrence on some plant species. Forthcoming approaches may gather important evidence to rank ants according to their capacity to defend plants from herbivores. We also advocate for established protocols to evaluate and compare ant aggressiveness and determine which species perform better than others.

## Acknowledgments

We are grateful to two reviewers and their comments that enhanced the quality of the manuscript. This work received no specific fund.

---

**Supplementary material 1.**
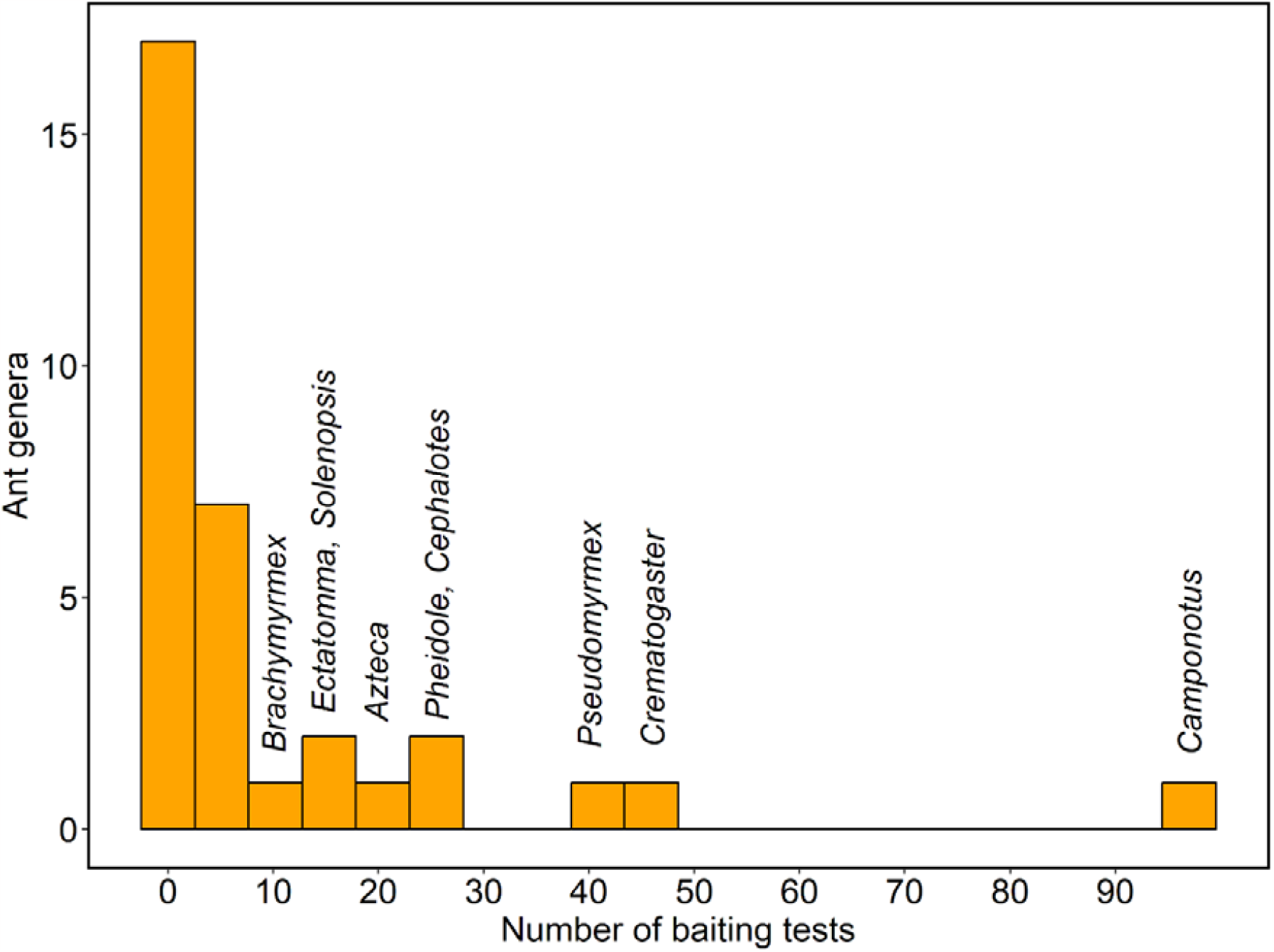
Histogram showing the frequency each ant studied in baiting studies. The first two columns refer to several other genera of ants. Data were retrieved from 56 published papers.

**Supplementary material 2.**
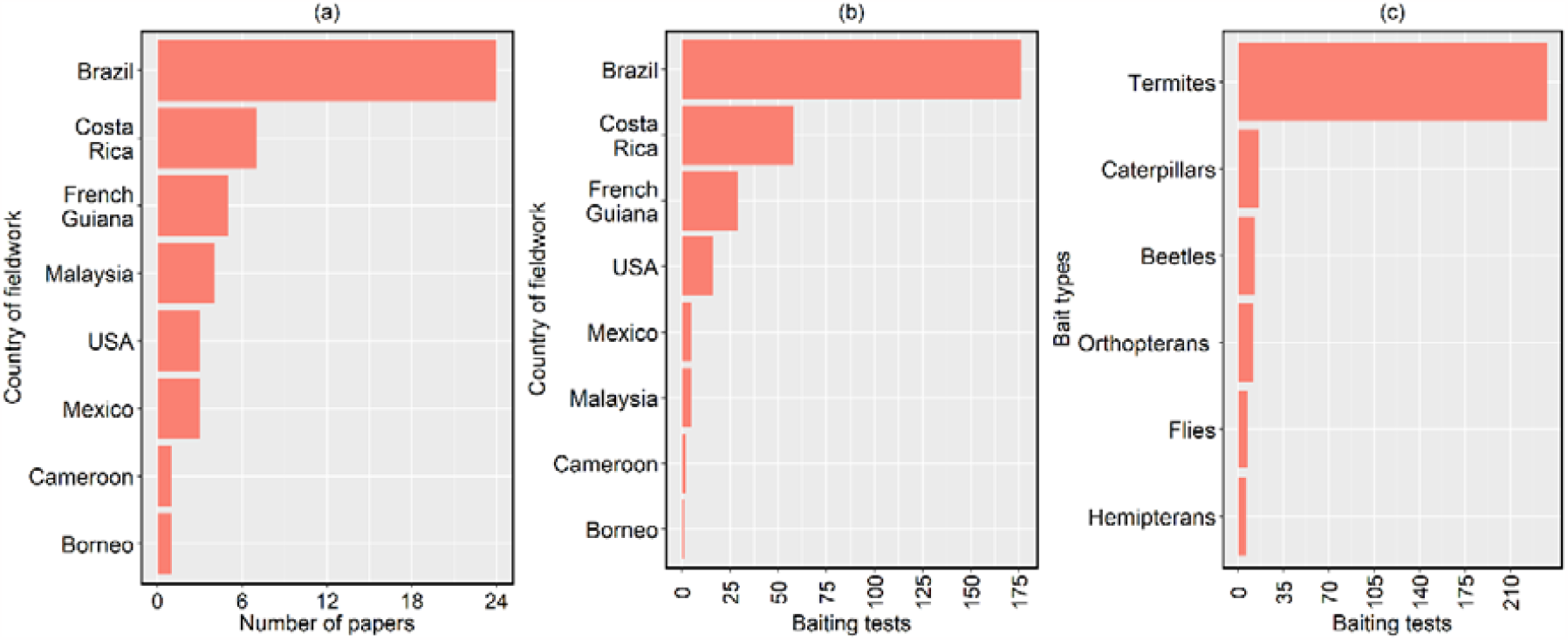
Overview of some characteristics of baiting studies that investigated ant aggressiveness. (a) Number of papers conducted in each country; (b) number of times ants were subjected to baiting tests in each country; (c) types of baits used in baiting tests. These data are not followed by statistical tests.

**Supplementary material 3.**
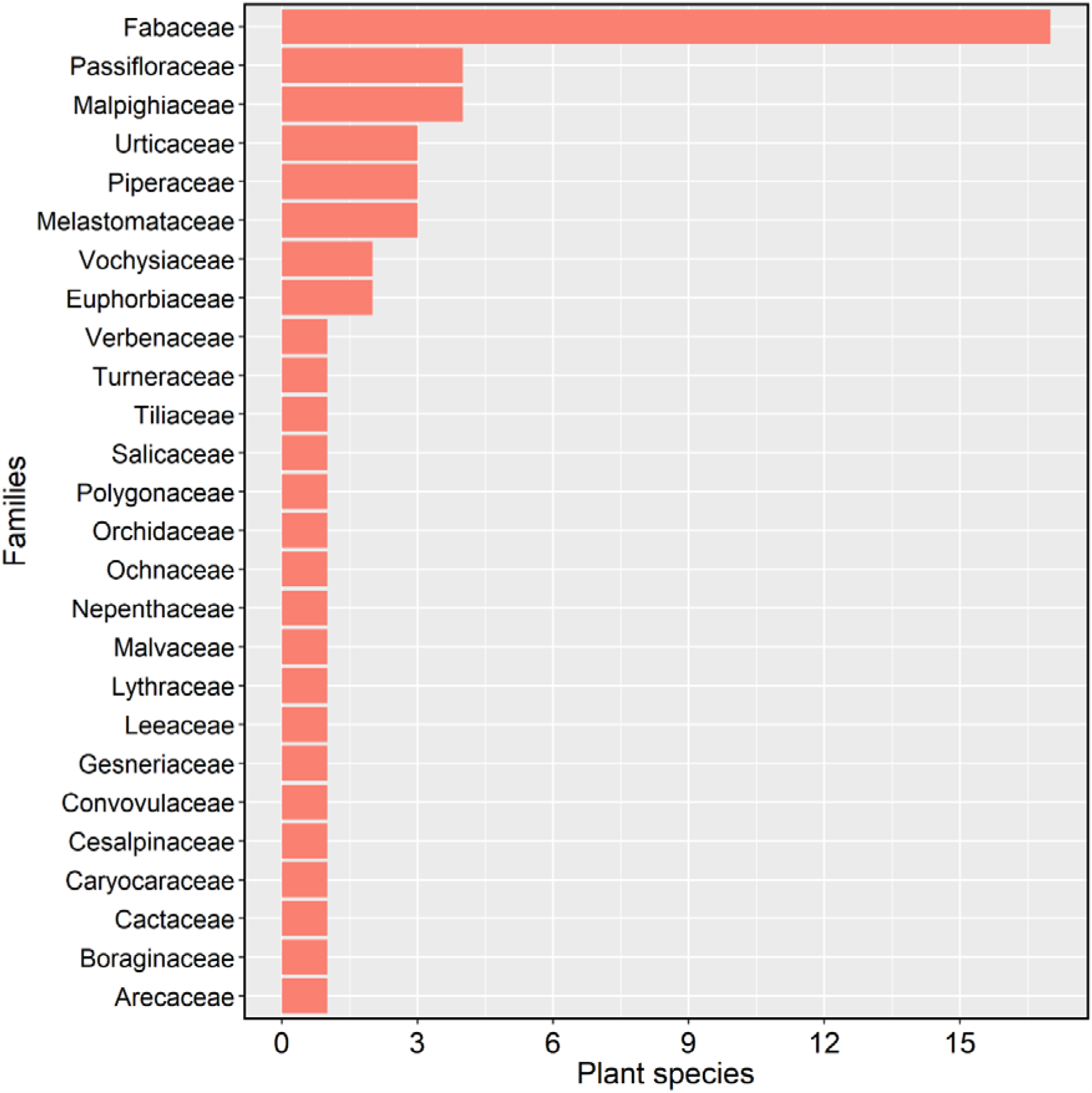
Frequency of species in each plant family, that was used in baiting tests.

**Supplementary material 4.**
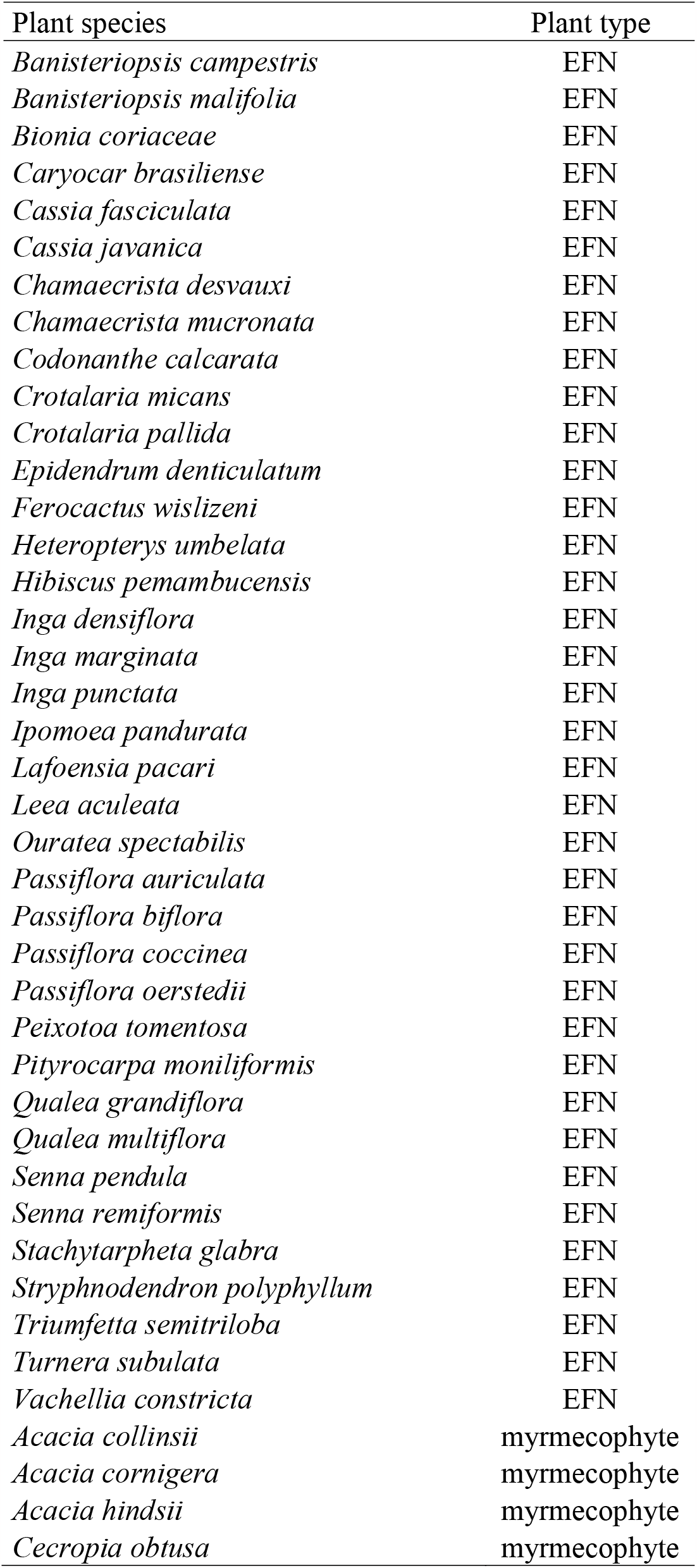
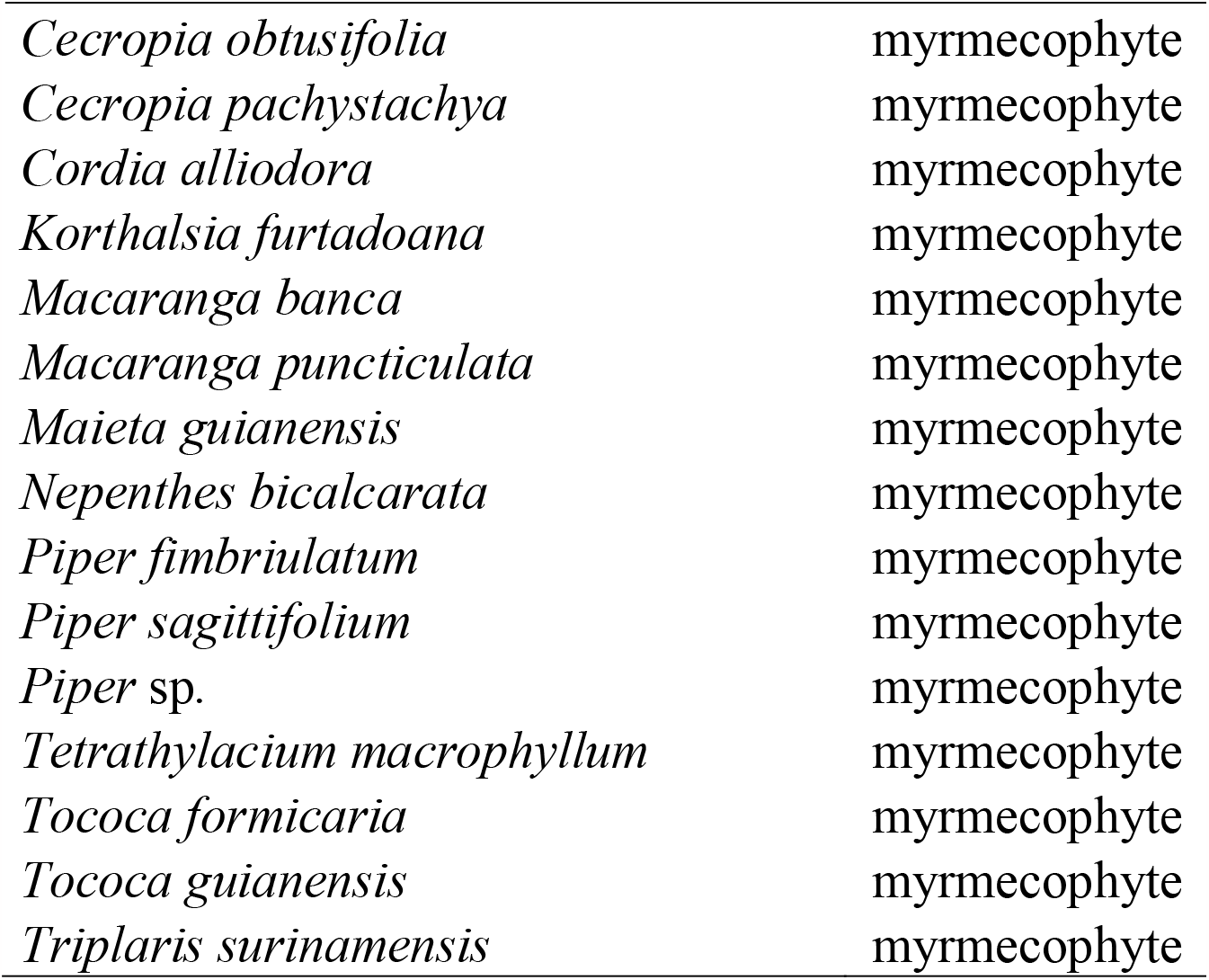
Plant species according to functional groups. EFN – plants with extrafloral nectaries.

| Plant species | Plant type |
| --- | --- |
| *Banisteriopsis campestris* | EFN |
| *Banisteriopsis malifolia* | EFN |
| *Bionia coriacea* | EFN |
| *Caryocar brasiliense* | EFN |
| *Cassia fasciculata* | EFN |
| *Cassia javanica* | EFN |
| *Chamaecrista desvauxi* | EFN |
| *Chamaecrista mucronata* | EFN |
| *Codonanthe calcarata* | EFN |
| *Crotalaria micans* | EFN |
| *Crotalaria pallida* | EFN |
| *Epidendrum denticulatum* | EFN |
| *Ferocactus wislizeni* | EFN |
| *Heteropterys umbelata* | EFN |
| *Hibiscus pemambucensis* | EFN |
| *Inga densiflora* | EFN |
| *Inga marginata* | EFN |
| *Inga punctata* | EFN |
| *Ipomoea pandurata* | EFN |
| *Lafoensia pacari* | EFN |
| *Leea aculeata* | EFN |
| *Ouratea spectabilis* | EFN |
| *Passiflora auriculata* | EFN |
| *Passiflora biflora* | EFN |
| *Passiflora coccinea* | EFN |
| *Passiflora oerstedii* | EFN |
| *Peixotoa tomentosa* | EFN |
| *Pityrocarpa moniliformis* | EFN |
| *Qualea grandiflora* | EFN |
| *Qualea multiflora* | EFN |
| *Senna pendula* | EFN |
| *Senna remiformis* | EFN |
| *Stachytarpheta glabra* | EFN |
| *Stryphnodendron polyphyllum* | EFN |
| *Triumfetta semitriloba* | EFN |
| *Turnera subulata* | EFN |
| *Vachellia constricta* | EFN |
| *Acacia collinsii* | myrmecophyte |
| *Acacia cornigera* | myrmecophyte |
| *Acacia hindsii* | myrmecophyte |
| *Cecropia obtusa* | myrmecophyte |
| *Cecropia obtusifolia* | myrmecophyte |
| *Cecropia pachystachya* | myrmecophyte |
| *Cordia alliodora* | myrmecophyte |
| *Korthalsia furtadoana* | myrmecophyte |
| *Macaranga banca* | myrmecophyte |
| *Macaranga puncticulata* | myrmecophyte |
| *Maieta guianensis* | myrmecophyte |
| *Nepenthes bicalcarata* | myrmecophyte |
| *Piper fimbriulatum* | myrmecophyte |
| *Piper sagittifolium* | myrmecophyte |
| *Piper* sp. | myrmecophyte |
| *Tetrathylacium macrophyllum* | myrmecophyte |
| *Tococa formicaria* | myrmecophyte |
| *Tococa guianensis* | myrmecophyte |
| *Triplaris surinamensis* | myrmecophyte |

